# Proofreading-deficient coronaviruses adapt over long-term passage for increased fidelity and fitness without reversion of exoribonuclease-inactivating mutations

**DOI:** 10.1101/175562

**Authors:** Kevin W. Graepel, Xiaotao Lu, James Brett Case, Nicole R. Sexton, Everett Clinton Smith, Mark R. Denison

**Affiliations:** Departments of Pathology, Microbiology and Immunology, Vanderbilt University Medical Center, Nashville, Tennessee, USA; Departments of Pathology, Microbiology and Pediatrics, Vanderbilt University Medical Center, Nashville, Tennessee, USA; Departments of Pathology, Microbiology and Elizabeth B. Lamb Center for Pediatric Research, Vanderbilt University Medical Center, Nashville, Tennessee, USA; Department of Biology, The University of the South, Sewanee, Tennessee, USA

## Abstract

The coronavirus (CoV) RNA genome is the largest among single-stranded positive sense RNA viruses. CoVs encode a proofreading 3′→5′exoribonuclease within nonstructural protein 14 (nsp14-ExoN) that is responsible for CoV high-fidelity replication. Alanine substitution of ExoN catalytic residues [ExoN(-)] in SARS-CoV and murine hepatitis virus (MHV) disrupts ExoN activity, yielding viable mutant viruses with defective replication, up to 20-fold decreased fidelity, and increased susceptibility to nucleoside analogs. To test the stability of the ExoN(-) genotype and phenotype, we passaged MHV-ExoN(-) 250 times in cultured cells (P250), in parallel with WT-MHV. Compared to MHV-ExoN(-) P3, MHV-ExoN(-) P250 demonstrated enhanced replication, reduced susceptibility to nucleoside analogs, and increased competitive fitness. However, passage did not select for complete or partial reversion at the ExoN-inactivating mutations. We identified novel amino acid changes within the RNA-dependent RNA polymerase (nsp12-RdRp) and nsp14 of MHV-ExoN(-) P250 that partially account for the observed changes in replication, susceptibility to nucleoside analogs, and competitive fitness observed in the passaged virus population, indicating that additional determinants can compensate for the activities of nsp14-ExoN. Our results suggest that while selection favors restoration of replication fidelity in ExoN(-) CoVs, there may be a significant barrier to ExoN(-) reversion. These results also support the hypothesis that high-fidelity replication is linked to CoV fitness and identify additional candidate proteins that may regulate CoV replication fidelity.

**IMPORTANCE:** Unique among RNA viruses, CoVs encode a proofreading exoribonuclease (ExoN) in nsp14 that mediates high-fidelity RNA genome replication. Proofreading-deficient CoVs with disrupted ExoN activity [ExoN(-)] are either non-viable or have significant defects in replication, RNA synthesis, fidelity, fitness, and virulence. In this study, we show that ExoN(-) murine hepatitis virus can adapt over long-term passage for increased replication and fitness without reverting the ExoN-inactivating mutations. Passage-adapted ExoN(-) mutants also demonstrate increasing resistance to nucleoside analogs that is only partially explained by secondary mutations in nsp12 and nsp14. These data suggest that enhanced resistance to nucleoside analogs is mediated by the interplay of multiple replicase proteins and support the proposed link between CoV fidelity and fitness.

## INTRODUCTION

A paradigm of RNA virus biology is error-prone genomic replication due to the lack of proofreading or post-replicative RNA repair mechanisms (1-3). Decreased replication fidelity may constrain RNA genome size and complexity and risks the accumulation of deleterious mutations leading to population extinction, a process known as lethal mutagenesis (4-7). While genetic diversity allows viral populations to adapt rapidly under selective pressure, many mutations are neutral or detrimental to viral fitness (8-12). Research performed with many RNA viruses supports the hypothesis that the mutation rate of RNA virus replicases has evolved to balance multiple characteristics of the viral population such as genetic diversity, genomic integrity, and virulence. High-or low-fidelity variants are described for many RNA viruses infecting animals including the coronaviruses (CoVs): murine hepatitis virus (MHV-A59) and Severe Acute Respiratory Syndrome-associated coronavirus (SARS-CoV) (13-17), as well as foot-and-mouth disease virus {Arias:2008bp, Zeng:2013dg, Zeng:2014dj, Xie:2014cs, Sierra:2007dg}, poliovirus (23-29), chikungunya virus (30, 31), influenza virus (32), coxsackievirus B3 (33, 34), and human enterovirus 71 (35-37). Most altered-fidelity variants described to date harbor mutations within the viral RNA-dependent RNA polymerase (RdRp), are attenuated *in vivo*, and protect against reinfection, highlighting their potential utility as live-attenuated vaccines (24, 28, 29, 38, 39). These studies underscore the importance of understanding the molecular mechanisms by which RNA viruses regulate their replication fidelity.

Viruses in the family *Coronaviridae* have large single-stranded positive-sense RNA genomes [(+)ssRNA] (40), ranging between 25.4 and 33.5 kilobases in length (41, 42). CoVs encode a 3′→5′ exoribonuclease (ExoN) in the N-terminal half of nonstructural protein 14 (nsp14-ExoN) (43, 44). CoV ExoN activity requires the presence of conserved magnesium-coordinating acidic amino acids in three motifs that together constitute the active site (DE-E-D) (45). The CoV ExoN is grouped with the DE-D-Dh superfamily of exonucleases involved in proofreading during prokaryotic and eukaryotic DNA replication (Fig. 1) (43-47). Alanine substitution of CoV motif I DE residues (DE→AA) reduces biochemical ExoN activity in SARS-CoV (45, 47) and human coronavirus 229E (43). MHV-A59 and SARS-CoV lacking ExoN activity [ExoN(-)] have mutation frequencies 8-to-20-fold greater than WT viruses (13, 14, 38). Thus, all available data to date support the hypothesis that nsp14-ExoN is the first known proofreading enzyme encoded by an RNA virus.

**Figure 1.**
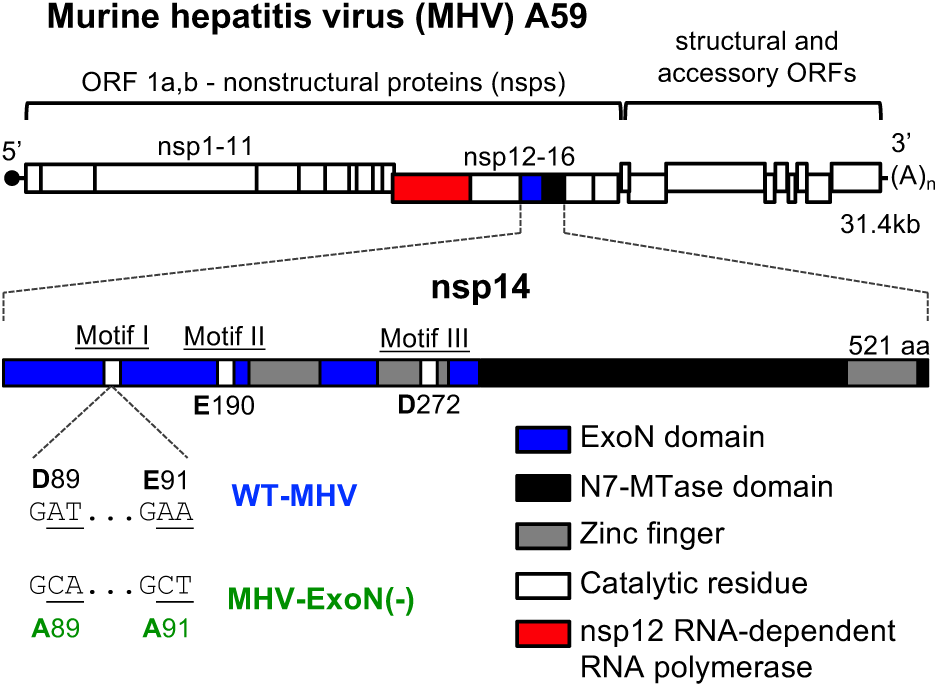
MHV genome organization and nsp14 exoribonuclease motifs. (*Top*) The MHV genome is a 31.4kb, capped (dark circle), and polyadenylated positive sense RNA molecule. The first two thirds of the genome encode 16 nonstructural proteins translated as a single polyprotein with a ribosomal frameshift. The final one third encodes the structural proteins. (*Inset*) Nsp14 encodes an exoribonuclease (solid blue) and N7-methyltransferase (hatched blue) and has 3 zinc fingers (gray boxes) predicted from the solved SARS nsp10/14 crystal structure (PDB 5C8U) (45). Catalytic residues for ExoN are marked with white boxes, and the engineered mutations for MHV-ExoN(-) are shown below the genome. The nsp12 RNA-dependent RNA polymerase is highlighted in red.

Despite the critical role of ExoN in virus replication, fidelity, fitness, and virulence, ExoN(-) mutants have not reverted the inactivating substitutions (G<u>AT</u>→G<u>CA</u> – D89A; G<u>AA</u>→G<u>CT</u> – E91A) (Fig. 1) (13, 14, 16, 17, 38). In fact, ExoN-inactivating mutations have been retained during up to 20 passages in culture, eight acute passages of SARS-CoV-ExoN(-) in aged BALB/c mice, and 60 days of persistent SARS-CoV-ExoN(-) infection in immunodeficient Rag-/-mice. In this study, we sought to determine whether long-term passage of MHV-A59-ExoN(-) (250 passages over one year; P250), hereafter MHV-ExoN(-), would result in virus extinction, ExoN(-) reversion, or compensation for the loss of proofreading. We demonstrate that MHV-ExoN(-) did not extinguish during the 250 passages and adapted in a gradual, continuous process for increased replication. MHV-ExoN(-) concurrently evolved reduced susceptibility to nucleoside and base analogs without reversion of the ExoN-inactivating mutations. The evolved mutations in MHV-ExoN(-) nsp14 and nsp12, which encodes the RdRp, accounted for only part of the increased replication and nucleoside analog resistance of MHV-ExoN(-) P250. This study demonstrates a surprising stability of the ExoN-inactivating substitutions, suggests that CoVs can evolve to compensate for the loss of ExoN functions, and indicates that proteins or other determinants outside of nsp12 and nsp14 likely contribute to CoV fidelity regulation.

## RESULTS

### Long-term passage of MHV-ExoN(-)

We serially passaged WT-MHV and MHV-ExoN(-) in delayed brain tumor (DBT) cells 250 times (P250). Virus from each passage was harvested once 50-100% of the monolayer was involved in syncytia, which occurred between 8 and 24 hours post-infection. Passage conditions varied for WT-MHV and MHV-ExoN(-) due to differences in replication kinetics between the two viruses. We stopped passage at P250 after observing reduced syncytia formation in MHV-ExoN(-)-infected flasks, likely resulting from a mutation in the MHV-ExoN(-) P250 spike protein cleavage site (discussed below).

### MHV-ExoN(-) and WT-MHV replicate with identical kinetics following 250 passages

MHV-ExoN(-) has a significant replication defect relative to WT-MHV (14). We first tested whether replication of MHV-ExoN(-) P250 was affected by long-term passage. MHV-ExoN(-) P3 replication was delayed by ∼2 hours and peak titer reduced by ∼1 log10 relative to WT-MHV P3 during high MOI infection (1 PFU/cell) (Fig. 2A, solid lines), consistent with our previous studies (14). By P250, both viruses replicated with identical kinetics (Fig. 2A, dotted lines). This represented a ∼1 log10 increase in peak replication for WT-MHV and a ∼2 log10 increase for MHV-ExoN(-), compared with the respective parental virus. Further, MHV-ExoN(-) P250 titers were maintained throughout the experiment, while WT-MHV P250 titers gradually decreased after 12 hours. We next performed infection at low MOI (0.01 PFU/cell) to examine replication over multiple cycles. Much like at MOI=1, WT-MHV P250 and MHV-ExoN(-) P250 replicated identically during multi-cycle infection, reaching peak titer around 28 hours post-infection (Fig. 2B). We also tested replication of MHV-ExoN(-) at P10, P50, P100, and P160. Replication kinetics gradually increased over passage, reaching P250-like levels by P100 (Fig. 2B). To determine whether the increased replication of MHV-ExoN(-) P250 was affected by the presence of potential defective viral genomes or by some other population-based phenomenon, both WT-MHV P250 and MHV-ExoN(-) P250 were plaque purified three times. The plaque-purified viruses replicated indistinguishably from the parent populations (Fig. 2C). Together, these data demonstrate that WT-MHV and MHV-ExoN(-) populations adapted for increased replication and that either individual genomes or those derived from a single virus plaque encoded the adaptive changes required by the total population.

**Figure 2.**
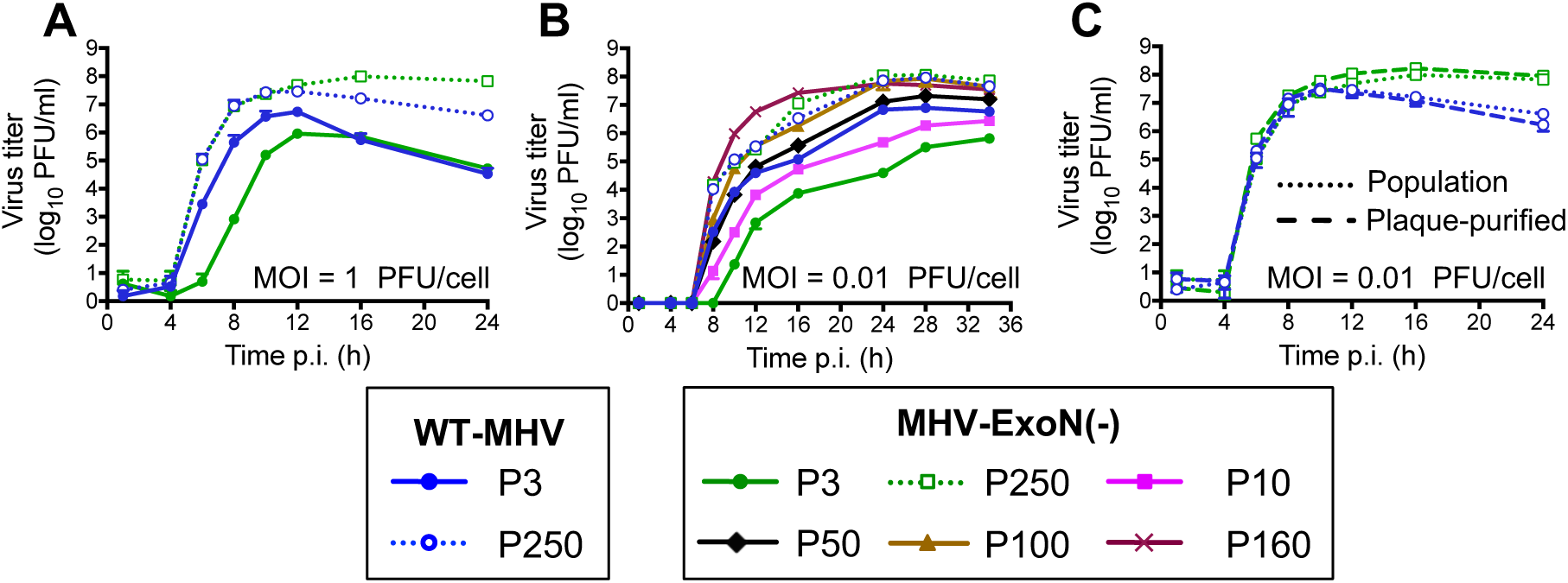
MHV-ExoN(-) evolved increased replicative capacity over long-term passage. Replication curves were performed for the indicated viruses during high MOI (1 PFU/cell) (*A*) and low MOI (0.01 PFU/cell) (*B*) infections. (*C*) WT-MHV P250 and MHV-ExoN(-) P250 were plaque purified three times, and replication curves were performed in parallel with the full population (MOI = 0.01 PFU/cell. Supernatants were collected at the indicated times post infection, and titers were determined by plaque assay. Data for *A-C* represent mean and standard deviation of n = 3.

### MHV-ExoN(-) accumulated 8-fold more mutations than WT-MHV but did not revert ExoN-inactivating mutations

To determine whether primary reversion of nsp14-ExoN(-) resulted in enhanced replication of MHV-ExoN(-) P250, we sequenced nsp14 from infected-cell total RNA. MHV-ExoN(-) P250 retained the motif I DE→AA substitutions, and we did not detect any variation in motif I at the level of di-deoxy sequencing at any passage, demonstrating that phenotypic changes were not the result of primary reversion of ExoN(-) motif I. To identify potentially adaptive consensus mutations, we performed full-genome di-deoxy sequencing of MHV-ExoN(-) P250 and WT-MHV P250. Within WT-MHV P250, we identified 23 mutations, of which 17 were nonsynonymous (NS) (Fig. 3A). In contrast, MHV-ExoN(-) P250 had 171 total mutations (74 NS) (Fig. 3B). The full-genome sequences have been deposited in GenBank, and the mutations for both viruses are listed in Supplemental Tables 1 and 2. We identified only one mutation shared by both viruses (nsp1 A146T), though it was present in approximately 50% of the WT-MHV P250 population by di-deoxy sequencing. Both viruses deleted most of the hemagglutinin esterase (HE). In MHV-A59, HE mRNA is not transcribed *in vitro* (48-50), and HE protein expression is detrimental to MHV-A59 fitness in cell culture (51). WT-MHV P250 also deleted ORF 4a, which is dispensable for MHV replication in cell culture (52). The C-terminal region of ns2 within MHV-ExoN(-) P250 was truncated and fused to HE with a -1 frameshift. Ns2 is a phosphodiesterase (PDE) that protects viral RNA by degrading 2′→5′ oligoadenylate, the activating factor for cellular RNase L (53-55). The portion of ns2 deleted in MHV-ExoN(-) P250 lies outside the PDE catalytic domain, in a region of unknown function. C-terminally truncated ns2 retains enzymatic activity (56), but whether this specific deletion and fusion disrupts PDE activity remains to be tested. Nevertheless, ns2 is dispensable for MHV replication in immortalized cells (57, 58). Details about the deletion sites are provided in Supplemental Figure 1. Within proteins predicted to be part of the replicase-transcriptase complex (nsp7-16 and nucleocapsid) (39), WT-MHV P250 had only one NS change, located in the nsp13-helicase (Fig. 3A). In contrast, MHV-ExoN(-) P250 had 17 NS changes within this region (Fig. 3B and Supplemental Tables 1, 2).

**Figure 3.**
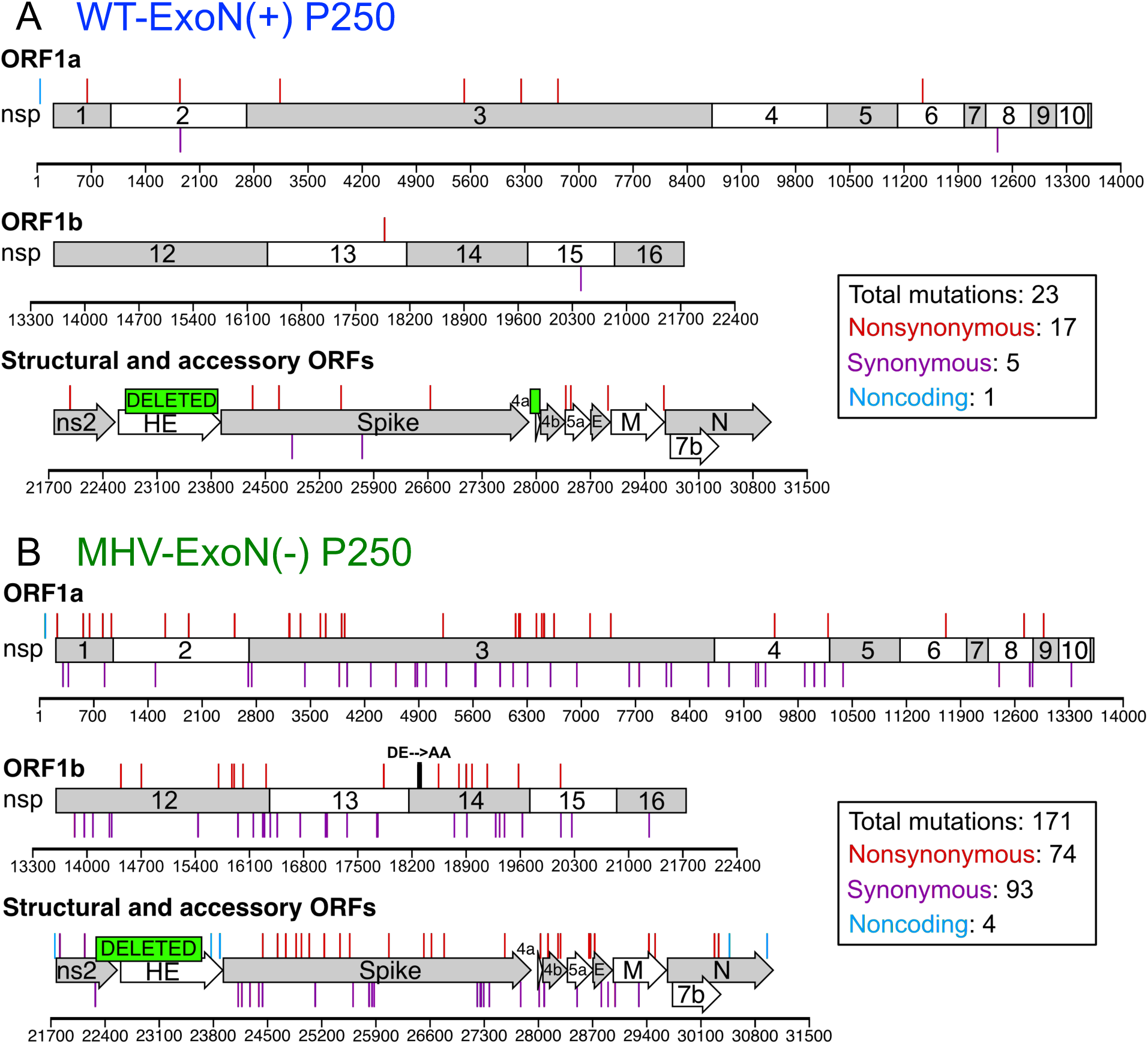
Mutation locations within passaged viruses. Mutations present at >50% by di-deoxy sequencing at passage 250 in WT-MHV (*A*) and MHV-ExoN(-) (*B*). Nonsynonymous mutations (red), noncoding mutations (cyan), and deletions (green boxes) are plotted above the schematic, and synonymous mutations (purple) are below.

### MHV-ExoN(-) P250 displays increased genomic RNA accumulation and increased resistance to 5-fluorouracil

Coronaviruses lacking ExoN consistently display defects in RNA synthesis relative to WT (14, 16, 43). To determine whether the increased replication of MHV-ExoN(-) P250 was associated with restored genomic RNA (gRNA) production, we measured gRNA accumulation over time using two-step real-time quantitative PCR (15, 16). MHV-ExoN(-) P250 accumulated similar levels of gRNA to WT-MHV P3 and WT-MHV P250 at early time points, while gRNA levels for MHV-ExoN(-) P3 were ∼1 log10 lower (Fig. 4A). MHV-ExoN(-) P250 gRNA levels fell below those of WT-MHV and WT-MHV P250 after 8 hours and were similar to those of MHV-ExoN(-) P3 at 10 hours post-infection. Normalizing to the gRNA abundance at four hours for each virus demonstrated that the rates of gRNA accumulation were similar for all four viruses (Fig. 4B). These data suggest that the increased replication of P250 viruses relative to WT-MHV is not fully accounted for by increased RNA synthesis. In addition to RNA synthesis defects, ExoN(-) CoVs have up-to 20-fold increased mutation frequencies and profoundly increased sensitivity to nucleoside and base analogs relative to WT CoVs (13, 14, 16, 17, 38). To determine whether nucleoside analog sensitivity of MHV-ExoN(-) was altered by long-term passage, we treated cells infected with parental and passaged viruses with the base analog, 5-fluorouracil (5-FU). 5-FU is converted intracellularly into a nucleoside analog that incorporates into growing RNA strands and causes A:G and U:C mutations. For simplicity, we hereafter refer to 5-FU as a nucleoside analog. Incorporation of 5-FU is increased in the absence of ExoN activity (16). All viruses displayed a concentration-dependent decrease in viral titer but differed greatly in their susceptibility to 5-FU (Fig. 4C). At 120μM, WT-MHV P3 titers were reduced by ∼1 log10, while MHV-ExoN(-) P3 titers were undetectable (> 5 log10-fold reduction). WT-MHV 5-FU sensitivity was not altered by passage. MHV-ExoN(-) P250 was less susceptible than MHV-ExoN(-) P3 to 5-FU treatment, with only a ∼1.5 log10 decrease in titer at 120 μM. MHV-ExoN(-) P250 remained more sensitive to 5-FU than WT-MHV, suggesting that WT-like resistance requires an intact ExoN. These data demonstrate that MHV-ExoN(-) P3 evolved resistance to 5-FU through mutations outside of ExoN(-) motif I.

**Figure 4.**
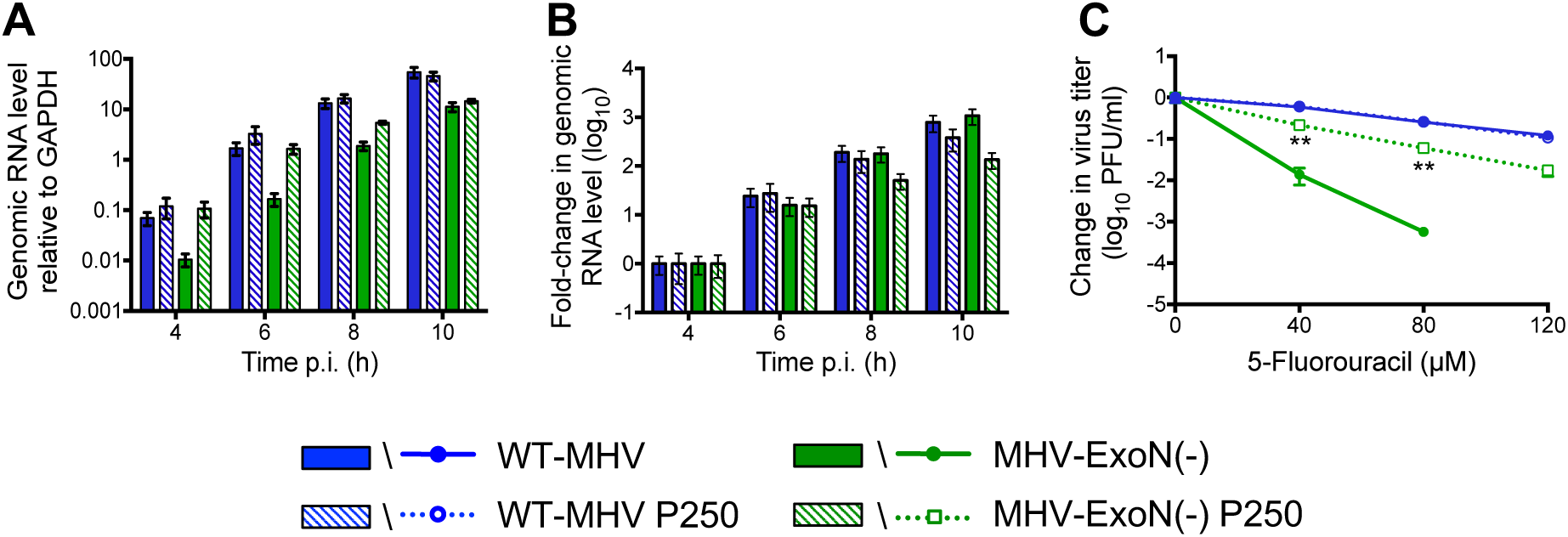
MHV-ExoN(-) P250 has increased genomic RNA accumulation and increased resistance to 5-fluorouracil. (*A*) Cells were infected with the indicated viruses at MOI = 1 PFU/cell, and intracellular RNA was harvested using TRIzol at the indicated times post infection. MHV genomic RNA was detected using SYBR green and primers directed to nsp10, and values were normalized to intracellular GAPDH. (*B*) Same data as in panel (*A*) normalized to RNA level for each virus at 4 hours post infection. Data represent mean and standard error for n = 9 (3 triplicate experiments). (*C*) Sensitivity of passaged viruses to 5-FU during low MOI infection (0.01 PFU/cell). Cells were treated with the indicated concentrations of 5-FU for 30 minutes prior to infection, and supernatants were harvested at 24 hours post-infection and titered by plaque assay. Data represent change in titer relative to untreated control and plotted as mean and standard error of n = 6 (two triplicate experiments). For panel C, statistical significance for change in titer of MHV-ExoN(-) P250 relative to MHV-ExoN(-) P3 was determined Mann-Whitney test (*P<0.05, **<0.01, ***P<0.001).

### Spike mutations in MHV-ExoN(-) P250 do not increase resistance to 5-FU

Bacteriophage ?X174 acquired resistance to 5-FU by delaying cell lysis, thereby reducing the number of replication cycles in which 5-FU can be incorporated (59). MHV-ExoN(-) P250 had multiple mutations in the spike glycoprotein, including one in the spike furin cleavage site that reduced syncytia formation. To test whether the spike mutations manifested in resistance to 5-FU, we cloned the spike gene from MHV-ExoN(-) P250 into the isogenic MHV-ExoN(-) background. The recombinant virus demonstrated intermediate replication kinetics between MHV-ExoN(-) P3 and MHV-ExoN(-) P250 (Fig. 5A) and did not form syncytia. Spike-P250 also increased the specific infectivity of viral particles (Fig. 5B). However, the MHV-ExoN(-) P250 spike did not affect sensitivity of the recombinant virus to 5-FU (Fig. 5C). Thus, any adaptive increase in 5-FU resistance must be located elsewhere in the genome.

**Figure 5.**
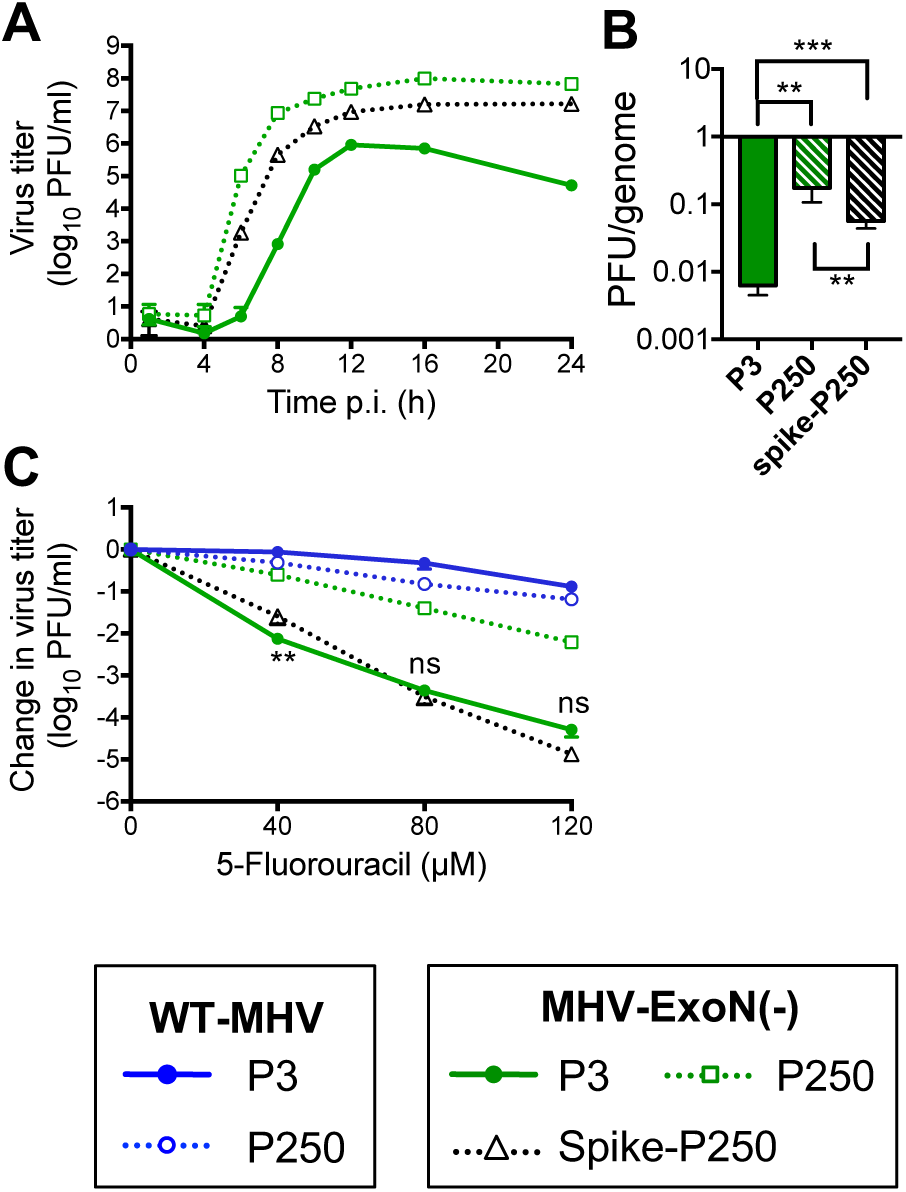
Mutations in the spike envelope protein from MHV-ExoN(-) P250 increase replicative capacity but do not affect sensitivity to 5-fluorouracil. (*A*) Replication kinetics of indicated viruses (MOI = 0.01 PFU/cell) plotted as mean and standard deviation with n = 3. (*B*) Specific infectivity of indicated viruses 12 hours post-infection (MOI = 1 PFU/cell). Data represent mean and standard error of n = 6 (two triplicate experiments). (*C*) Sensitivity of indicated viruses to 5-fluorouracil during low MOI infection (0.01 PFU/cell), as described in Fig. 4. Data represent mean and standard error of n = 6 (two triplicate experiments). For panel B, statistical significance was determined using one-way ANOVA. For Panel C, statistical significance for change in titer of MHV-ExoN(-) spike-P250 relative to MHV-ExoN(-) P3 was determined using Mann-Whitney test (*P<0.05, **<0.01, ***P<0.001, ns = not significant).

### MHV-ExoN(-) passage resulted in unique mutations in nsp12 and nsp14

To date, three proteins have been shown to alter CoV sensitivity to 5-FU: the nsp12-RdRp, nsp14-ExoN, and nsp10, which stimulates ExoN activity (15, 17, 39). Neither WT-MHV nor MHV-ExoN(-) P250 contained a NS mutation in nsp10, and WT-MHV P250 had no mutations within either nsp12 or nsp14. In contrast, MHV-ExoN(-) P250 had 7 NS mutations in nsp12 and 6 NS mutations in nsp14 (Figs. 3, 6), none of which have been described previously *in vitro* or in viable viruses. Within nsp12, six mutations were in the predicted RdRp fingers, palm, and thumb domains (Fig. 6A) (60). Four residues (H709, F766, S776, and M814) can be visualized on a Phyre^2^-modeled structure of the MHV-nsp12 RdRp, while the remainder lie outside the modeled core RdRp (Fig. 6A) (17). One mutation, M288T, lies in the CoV-specific domain, which is conserved among nidoviruses. This domain has been implicated in membrane targeting in MHV-A59 (61) and performs an essential nucleotidylation activity in the arterivirus, equine arteritis virus (62). However, M288T is not predicted to catalyze nucleotidylation. Within nsp14, 4 NS mutations were identified in the ExoN domain, and 2 NS mutations were in the C-terminal N7-methyltransferase domain (Fig. 6B). We next modeled the structure of MHV nsp14 using Phyre^2^ software (63), resulting in highest-probability similarity to the SARS-CoV nsp14-nsp10 complex (PDB: 5C8S) (45) with high-confidence (i.e. the calculated probability of true homology between the structures) of 100% for residues 3-519 of MHV-nsp14. The model predicts that five mutations are located close to surface of the protein (Fig. 6B). All three modeled zinc finger domains contain one NS mutation (F216Y, Y248H, L473I). Two mutations, D128E and F216Y, are located near the interface between nsp10 and nsp14, though neither site has previously been implicated in nsp10-nsp14 interaction (15, 64, 65). One NS mutation resulted in a D272E substitution in ExoN motif III, a metal-coordinating active site residue. We previously reported that alanine substitution of D272 results in an ExoN(-) phenotype (14), but the viability or phenotype of a D272E substitution was not tested in that study. These data suggest that a network of residues evolved to regulate nsp12 and nsp14 activity or stability in the ExoN(-) background.

**Figure 6.**
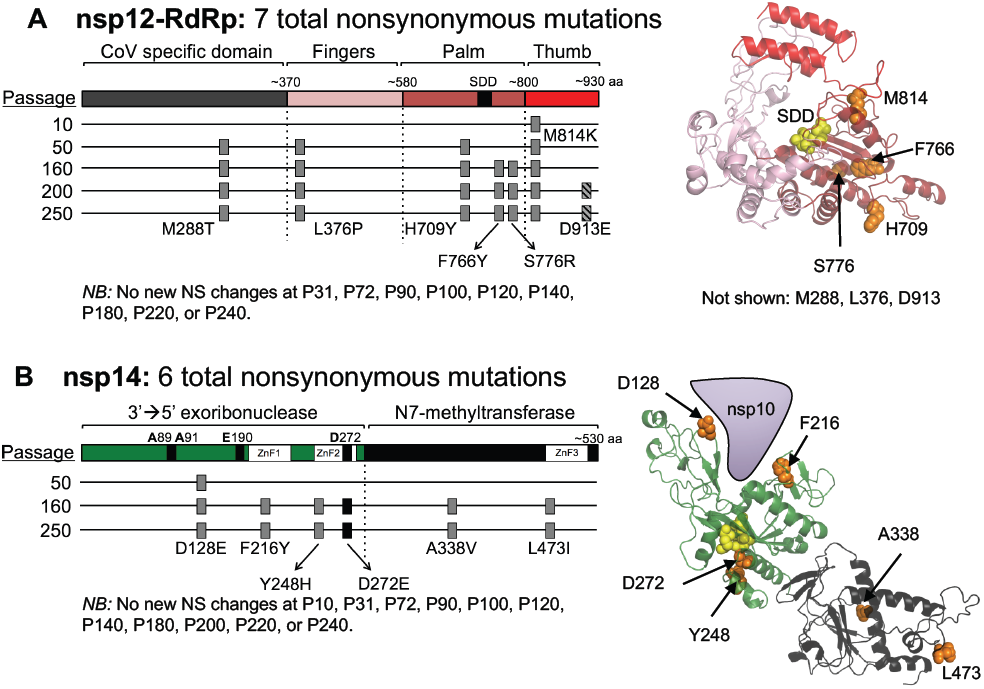
The timing of fixation of mutations in nsp12-RdRp and nsp14-ExoN within MHV-ExoN(-) correlates with increased resistance to nucleoside analogs. (*A*) A schematic of nsp12-RdRp with the CoV-specific region and the canonical fingers, palm, and thumb domains of RdRps is shown. The nsp12-RdRp coding region was sequenced at the indicated passage, and the nonsynonymous changes are plotted; gray boxes indicate consensus changes, and hatched boxes indicate variants present at <50% of the population by di-deoxy sequencing. At right, mutations are marked in orange on a Phyre^2^-modeled structure of MHV-nsp12, with the active site residues marked in yellow (17). RdRp domains are colored according to the linear schematic. *NB*: M288T, L376P, and D913E lie outside the modeled region and thus are not marked. (*B*) A schematic of nsp14 with the ExoN and N7-methyltransferase domains is shown, with mutation plotting as in panel A. The black box denotes a mutation to ExoN motif III. At right, mutations are marked in orange on a Phyre^2^-modeled structure of the MHV-nsp14/nsp10 complex. Domains are colored according to the linear schematic.

### Fixed mutations in nsp12 and nsp14 in MHV-ExoN(-) P250 directly correlate with increased resistance to multiple nucleoside analogs

To determine approximately when the mutations in nsp12 and nsp14 arose, we performed di-deoxy sequencing across these protein-coding regions roughly every 20 passages (P10, 31, 50, 72, 90, 100, 120, 140, 160, 180, 200, 220, 240). By this method, we detected consensus NS mutations at P10, P50, and P160 for nsp12, and at P50 and P160 for nsp14 (Fig. 6). Both nsp12 and nsp14 carried their full complement of P250 consensus mutations by P160, except for a minority variant (D913E) in nsp12 maintained at <50% of the population between P200 and P250. These passage levels correlated with increased replication of MHV-ExoN(-) (Fig. 2B) and with decreasing sensitivity to 5-FU (Fig. 7A). Neither replication nor 5-FU sensitivity of MHV-ExoN(-) changed substantially between P160 and P250. To determine whether MHV-ExoN(-) evolved increased resistance to multiple nucleoside analogs, we treated virus-infected cells with three additional analogs that are substrates for viral RdRps: ribavirin (RBV), a guanine analog that inhibits viral replication through multiple mechanisms, including mutagenesis and inhibition of purine biosynthesis (66); 5-azacytidine (AZC), an RNA mutagen (67); and 2’-C-methyladenosine (CMeA), which is proposed to incorporate in viral RNA and terminate nascent transcripts (68). As with 5-FU, we observed dose-dependent sensitivity to RBV, AZC, and CMeA in all MHV-ExoN(-) viruses that decreased with increasing passage number (Fig. 7B-D). Except against AZC, MHV-ExoN(-) sensitivity did not change between P160 and P250. Together, these data demonstrate that MHV-ExoN(-) evolved increased resistance to multiple nucleoside analogs that correlated with the length of passage and the acquisition of mutations in nsp12 and nsp14. Importantly, this occurred in the absence of specific mutagenic selection and without reversion of ExoN motif I. This increased general selectivity towards all four classes of nucleotide strongly advocates for an overall increase in fidelity in MHV-ExoN(-) P250.

**Figure 7.**
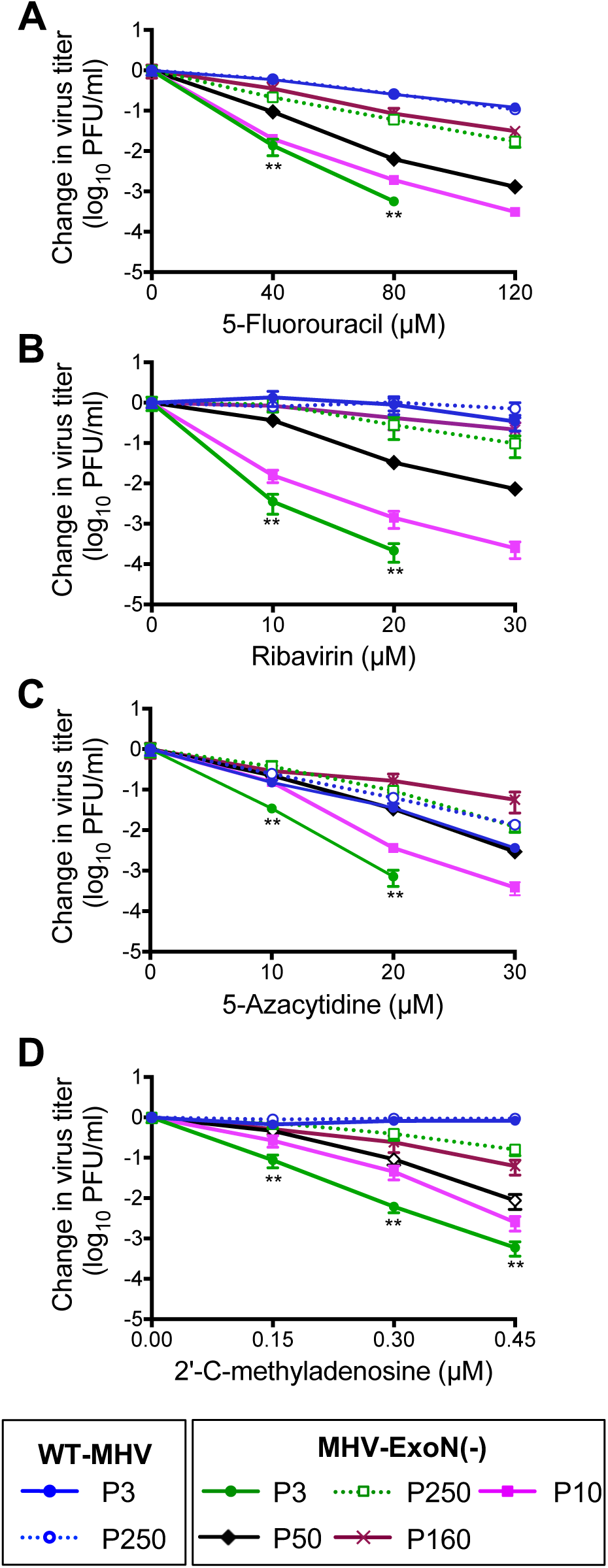
MHV-ExoN(-) evolved increased resistance to nucleoside and base analogs over passage. Sensitivity of isogenic and passaged viruses to the nucleoside analogs, 5-FU (*A*), ribavirin (*B*), 5-azacytidine (*C*), and 2’-C-methyladenosine (*D*) during low MOI infection (0.01 PFU/cell) as described in Fig. 4. Data represent mean and standard error for n = 6 (2 triplicate experiments). *NB*: Data for WT-MHV, WT-MHV P250, MHV-ExoN(-) P3, MHV-ExoN(-) P250 in panel (*A*) are the same as shown in Fig. 4C. Statistical significance for change in titer of MHV-ExoN(-) P3 relative to MHV-ExoN(-) P250 was determined using Mann-Whitney test (*P<0.05, **<0.01, ***P<0.001).

### Mutations in nsp12 partially account for increased resistance of MHV-ExoN(-) P250 to multiple nucleoside analogs

We hypothesized that mutations in MHV-ExoN(-) P250 nsp12 and nsp14 were most likely to impact replication and nucleoside analog sensitivity based on their enzymatic activities and temporal association with phenotypic changes. To test this hypothesis, we engineered recombinant MHV-ExoN(-) to encode the P250 nsp12 and nsp14, alone and together. Expression of nsp12-P250 and nsp14-P250, alone or in combination, altered replication kinetics of MHV-ExoN(-) without affecting peak titers (Fig. 8A) and increased gRNA levels above those of MHV-ExoN(-) P3 (Fig. 8B). Nsp12-P250 had a greater effect than nsp14-P250 on the sensitivity of MHV-ExoN(-) to all analogs tested, and the combination of nsp12-and nsp14-P250 did not increase resistance above nsp12-P250 alone (Fig. 8C-E). None of the recombinant viruses recapitulated the resistance phenotypes of the MHV-ExoN(-) P250 population. Together, these data demonstrate that nsp12-P250 mutations only partially account for the nucleoside analog resistance of MHV-ExoN(-) P250, and that adaptations in nsp12-P250 mask those in nsp14-P250. We also can conclude that the nsp14-P250 D272E active site mutation does not correct the defect caused by the motif I DE→AA substitutions. Together, the results suggest possible limitations for fidelity adaptation in nsp12 and nsp14, and also the exciting prospect that other CoV proteins participate in fidelity regulation within MHV-ExoN(-) P250.

**Figure 8.**
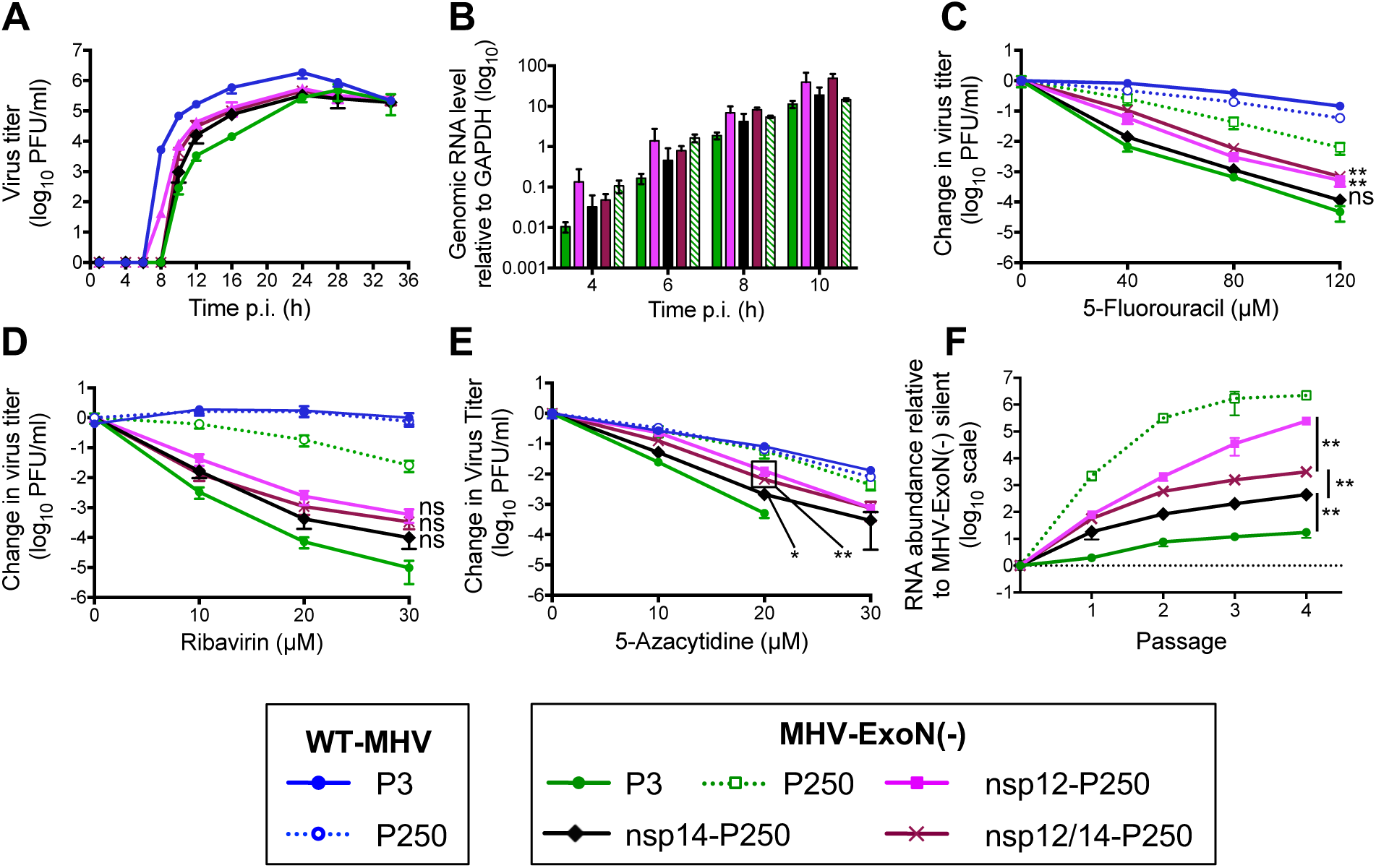
Mutations in nsp12-RdRp and nsp14-ExoN from MHV-ExoN(-) P250 incompletely increase resistance to RNA mutagens and increase fitness of MHV-ExoN(-). (*A*) Replication kinetics of recombinant P250 viruses (MOI = 0.01 PFU/cell) plotted as mean and standard deviation with n = 3. (*B*) Genomic RNA accumulation relative to intracellular GAPDH, as described in Fig. 4. Data represent mean and standard error for n = 6-9 (2-3 triplicate experiments). (*C-E*) Sensitivity of recombinant MHV-ExoN(-) viruses to 5-FU (*C*), ribavirin (*D*), and 5-azacytidine (*E*) during low MOI infection (0.01 PFU/cell) as described in Fig. 4. Data represent mean and standard error of n = 6. (*F*) Recombinant viruses were competed against a reference MHV-ExoN(-) containing 10 silent mutations within nsp2. The ratio of viral genome copies relative to the MHV-ExoN(-) reference is plotted. Data represent mean and standard error of n = 6. *NB*: MHV-ExoN(-) P250 data set contains 4 replicates at passage 3 and a single replicate at passage 4 due to undetectable levels of MHV-ExoN(-) silent. For panels C-E, statistical significance for change in titer of swapped viruses relative to MHV-ExoN(-) P3 at highest drug concentration tolerated was determined using Mann-Whitney test (*P<0.05, **<0.01, ***P<0.001, ns = not significant). For panel F, statistical significance for the indicated comparisons was determined using Mann-Whitney test. Boxed points have the same P value.

### Resistance to nucleoside analogs correlates with MHV-ExoN(-) fitness

We hypothesized that mutations in nsp12 and nsp14 provided a fitness advantage to MHV-ExoN(-) P250. We competed the recombinant viruses with a reference MHV-ExoN(-) virus (P1 stock) containing 10 silent mutations in the nsp2 coding region. Mutant and reference viruses were detected in the mixed infection by real-time quantitative PCR using dual-labeled probes specific for each virus. MHV-ExoN(-) P3 showed a modest fitness advantage over the reference P1 MHV-ExoN(-) silent (Fig. 8F, solid green). MHV-ExoN(-) P250 profoundly outcompeted MHV-ExoN(-) silent, with >1000-fold more MHV-ExoN(-) P250 genomes present at the end of passage 1 (Fig. 8F, dotted green). MHV-ExoN(-) nsp12-P250 had greater relative fitness than MHV-ExoN(-) nsp14-P250, and MHV-ExoN(-) nsp12/14-P250 was intermediate between the single recombinants, implicating a complex evolutionary interaction between these two proteins. The measured fitness correlated with the patterns of nucleoside analog resistance and RNA synthesis associated with mutations in nsp12 and nsp14, suggesting a link between the evolution of these virus phenotypes.

The result also confirms that nsp12 and nsp14 are important but not sufficient to account for the significantly increased fitness of MHV-ExoN(-) P250 relative to MHV-ExoN(-) P3.

## DISCUSSION

High-fidelity replication is proposed to be critical for maintaining the large (+)ssRNA genomes of CoVs (40, 69). Our previous studies demonstrate that ExoN-mediated proofreading is required for replicative fitness, nucleoside analog resistance, and virulence (16, 17, 38). Yet no direct test of the requirement for ExoN activity for long-term survival and stability of the CoV genome had been performed. In this study, we describe experimental adaptive evolution of WT-MHV and MHV-ExoN(-) during long-term passage in cell culture. WT-MHV evolved increased replication kinetics over 250 passages, with few consensus mutations arising in the WT-MHV P250 genome. In contrast, MHV-ExoN(-) accumulated 8-fold more mutations than WT-MHV, none of which occurred at the ExoN-inactivating mutations. Nevertheless, MHV-ExoN(-) P250 demonstrated increased replication kinetics and increased resistance to nucleoside analogs as compared to MHV-ExoN(-) P3. Mutations within nsp12-P250 conferred greater resistance to nucleoside analogs than nsp14-P250, suggesting that secondary mutations within nsp14 cannot restore ExoN activity without the full complement of catalytic DE-E-D residues. These results further support our studies and those from other exonucleases that the motif I DE residues are absolutely required for ExoN activity. However, mutations in nsp12-P250 also were not sufficient, alone or in combination with nsp14-P250, to account for the increased nucleoside analog resistance of the MHV-ExoN(-) P250 virus population, indicating that other proteins contribute to resistance or that the virus evolved increased mutational robustness over passage.

### Why doesn’t MHV-ExoN(-) revert over long-term passage?

We detected no primary reversion at the DE→AA substitutions MHV-ExoN(-) at any passage tested. These data are consistent with and significantly extend previous studies reporting genotypic stability of ExoN(-) motif I in MHV and SARS-CoV (13, 14, 16, 17, 38). Complete reversion within ExoN(-) motif I to DE would require four nucleotide changes. This likely represents a high genetic barrier to reversion, especially given that fitness can be increased by mutations outside of nsp14-ExoN. Single and double nucleotide changes within motif I could restore acidic charge to individual residues (e.g. motif I EA, AD, ED, etc). However, the active site compositions of DEDDh exonucleases, such as the Klenow fragment, are so stringent that even conservative mutations (D-to-E or E-to-D) reduce ExoN activity by >96% (70). Thus, intermediate amino acid changes may not have a selective advantage compared to motif I AA, limiting the evolutionary pathways to reversion. However, nsp14-P250 increased competitive fitness in the MHV-ExoN(-) background (Fig. 8F), demonstrating a modest capacity for fitness adaptation in nsp14 outside of the catalytic residues. Whether these mutations resulted from genetic drift or positive selection remains unclear. Nevertheless, our data show that MHV-ExoN(-) can adapt for increased fitness without fully restoring exoribonuclease activity. Understanding the mechanisms by which MHV-ExoN(-) P250 compensated for ExoN activity could allow recovery of ExoN(-) variants of other CoVs, such as transmissible gastroenteritis virus and human CoV 229E, which to date have been unrecoverable as ExoN(-) recombinants (43, 71).

### CoV adaptation to loss of ExoN-mediated proofreading

High-fidelity replication, made possible in part by the proofreading activity of nsp14-ExoN, is thought to have been essential for the expansion and maintenance of large CoV genomes (39, 40, 69). Compensation for the loss of ExoN could have occurred through the evolution of increased fidelity via other replicase proteins. Mutations conferring increased replication fidelity to RNA viruses have most frequently been mapped to RdRps (24, 25, 30, 72). Three findings suggest that mutations within nsp12-P250 confer increased polymerase fidelity. First, nonsynonymous mutations to the RdRp arose in the low-fidelity MHV-ExoN(-) but not in the presence of proofreading (WT-MHV). Second, five of the mutations lie in or near structural motifs important for fidelity regulation in other RdRps.

Amino acid substitutions in the fingers and palm domains have been repeatedly shown to affect viral RdRp fidelity (25, 34), and we have recently reported a fingers mutation (nsp12-V553I) that likely increases the fidelity of the MHV RdRp (17). Our modeled structure predicts that nsp12-P250 contains three mutations in the palm domain and one in the fingers domain, with the M814K thumb domain mutation lying near the palm (Fig. 6A). Third, exchange of nsp12-P250 alone into the background of MHV-ExoN(-) reduced the susceptibility of MHV-ExoN(-) to three different nucleoside analogs (Fig. 8). Although resistance to a single nucleoside analog can evolve without increasing overall fidelity, resistance to multiple nucleoside analogs suggests a broadly increased capacity to discriminate modified nucleotides {Arias:2008bp, Sierra:2007dg, Zeng:2013dg, Guo:2016fx}. Thus, nsp12-P250 is likely a high-fidelity polymerase. However, nsp12-P250 only partially accounts for the MHV-ExoN(-) P250 nucleoside analog resistance phenotype (Fig. 8), suggesting a possible limit to the compensation achievable by mutating the RdRp alone. Further, the effects of mutations in nsp12-P250 and nsp14-P250 are not additive and may be antagonistic when isolated from the whole passaged virus (Fig. 8), indicating that the relationships between nsp12-and nsp14-P250 mutations are likely evolutionarily linked with those in other MHV proteins. In fact, a substantial component of the evolved resistance to nucleoside analogs cannot be explained by nsp12-P250 and nsp14-P250, alone or together. In support of this hypothesis, we identified several nonsynonymous mutations in other replicase proteins, such as nsps 8, 9, 13, and 15. SARS-CoV nsp8 and nsp13 have functional interactions with nsp12, acting as a primase/processivity factor (74, 75) and a helicase/NTPase, respectively (76). Processivity factors in herpes simplex virus and *Mycobacterium tuberculosis* regulate DNA polymerase fidelity by balancing polymerase extension and exonuclease activity (77, 78), and helicases in chikungunya virus and foot-and-mouth disease virus can evolve to increase fidelity (79) and alter the frequency of ribavirin-induced mutations (80), respectively. SARS-CoV nsp9 has RNA-binding activities and is proposed to participate in the multi-protein replicase complex (39, 81), and MHV nsp15 is a uridylate-specific endoribonuclease (82). Both could plausibly be involved in modifying polymerase activity. Additionally, it remains possible that evolution for increased fidelity could involve proteins outside the canonical replication complex (nsps7-16), including those in the structural and accessory cassette. Thus, while immediate studies will focus on testing whether replicase proteins nsp8, 9, 13, and 15 regulate fidelity, it is exciting to consider the possibility that this virus-directed discovery approach will reveal novel interactions between multiple MHV proteins.

### Has MHV-ExoN(-) P250 evolved robustness as a defense against ExoN(-) infidelity?

Three of the nucleoside analogs used in this study (5-FU, ribavirin, and 5-azacytidine) reduce virus titers at least in part through mutagenesis (16, 66, 67). Although the resistance of MHV-ExoN(-) P250 to these RNA mutagens may be fully explained by the evolution of a high-fidelity replication complex, an additional possibility is that MHV-ExoN(-) P3 evolved increased mutational robustness. Mutational robustness is the capacity of a virus to buffer the fitness effects of mutations, which are most frequently detrimental or lethal (8-12). Thus, in a low-fidelity virus like MHV-ExoN(-) P3, genomic mutations which reduce the fitness cost of subsequent mutations can provide a selective advantage (83-85). The relationship between fidelity and robustness has been demonstrated for poliovirus and coxsackievirus B3: wild-type poliovirus encodes a lower-fidelity polymerase than coxsackievirus and is more mutationally robust (86). Selection for increased robustness could explain the ∼90 synonymous changes in MHV-ExoN(-) P250. Synonymous changes can alter codons to reduce the probability of non-conservative amino acid changes (87). Alternatively, the increased population size of MHV-ExoN(-) P250 could promote robustness by a “safety-in-numbers” effect, allowing efficient purging of low-fitness mutants while maintaining population fitness (88). Large populations also increase the likelihood of co-infection, allowing complementation between viral genomes. We did not observe an effect of increased replication in MHV-ExoN(-) P250 nucleoside analog resistance (Fig. 5), but a recent study with poliovirus suggests that mutagenized populations have elevated coinfection frequencies (89). Thus, complementation may contribute to MHV-ExoN(-) P250 nucleoside analog resistance. Conflicting evidence exists regarding whether mutational robustness itself affects the sensitivity to RNA mutagens (86, 87, 90); nevertheless, the robustness of MHV-ExoN(-) P250 merits further investigation.

### Conclusions

One of the defining features of CoV replication is the proofreading activity of the nsp14 exoribonuclease that is a critical determinant of CoV replication, fidelity, and fitness. We here show that CoVs also have the capacity to compensate for loss of ExoN activity through a network of mutations in nsp12, nsp14, and elsewhere in the genome. Thus, while nsp14-ExoN appears to play a dominant role in CoV replication fidelity, its activity is likely closely tied to a highly evolved network of proteins. The demonstrated co-adaptation for replication, fidelity, and competitive fitness within MHV-ExoN(-) supports the hypothesis that these roles are linked functionally and evolutionarily. Genetic and biochemical testing of the rich mutational resource revealed in MHV-ExoN(-) P250 will likely inform the design of countermeasures for endemic and emerging CoVs by defining novel common targets for stable virus attenuation or direct inhibition.

## MATERIALS AND METHODS

### Cell culture

DBT-9 (delayed brain tumor, murine astrocytoma clone 9) cells were maintained as described previously (91). Baby hamster kidney (BHK) cells stably expressing the MHV-A59 receptor, CEACAM1, [BHK-R, (15)] were maintained under selection with 0.8 mg/ml of G418 (Mediatech) as described previously (91).

### Long-term passage of virus and stock generation

The infectious cDNA clone for MHV-A59 and the recovery of MHV-ExoN(-) are described previously (14, 91). Long-term passage was initiated by infecting subconfluent monolayers of DBT-9 cells in 25 cm2 flasks with either wild-type MHV-A59 or MHV-ExoN(-) at a multiplicity of infection (MOI) of approximately 0.1 particle forming unit per cell (PFU/cell). One lineage of each virus was passaged blindly for a total of 250 passages (P250). Supernatant was harvested at each passage and stored at -80°C. Total RNA was harvested for most passages using 1 ml of TRIzol Reagent (Ambion) per 25 cm2 flask and stored at -80°C. Virus stocks of select intermediate passages were generated by infecting a subconfluent 150 cm2 flask of DBT-9 cells at an MOI of 0.01 PFU/cell.

Approximately 24 hours post-infection the flask was frozen at -80°C and the supernatant was clarified by centrifugation at 4,000 x g (Sorvall RC 3B Plus; HA-6000A rotor) for 10 min at 4°C. The virus titer of each stock was determined by plaque assay using DBT-9 cells as described previously (14, 91). For plaque assays of viruses containing the spike protein from MHV-ExoN(-) P250, which does not form syncytia, plaques were visualized with neutral red (Sigma #N6264, diluted 1:10 in PBS containing calcium and magnesium). Neutral red was added 24 hours after plating and incubated for an additional 3-8 hours before formaldehyde fixation. Plaque purification was performed by infecting DBT cells with serial dilutions of virus and overlaying the cultures with agar. Single plaques were isolated, resuspended in PBS containing calcium and magnesium, and inoculated onto fresh DBTs. This process was completed 3 times before experimental stocks were generated, as above.

### Sequencing of virus stocks

Following P250, RNA was purified from the harvested TRIzol samples according to the manufacturer’s protocol and reverse transcribed (RT) using SuperScript III (Invitrogen) as described previously (14). Full-genome di-deoxy sequencing was performed for both WT-MHV P250 and MHV-ExoN(-) P250 using 12 overlapping amplicons approximately three kilobases in length. All coding regions were sequenced fully, and out of 31,409 nucleotides, > 99% were sequenced for each virus [WT-MHV P250: 21-31,279 and MHV-ExoN(-) P250: 21-31,275]. Two microliters of RT product were used for each PCR reaction (16). Di-deoxy sequencing was performed by Genhunter Corporation (Nashville, TN) or GENEWIZ (South Plainfield, NJ). Sequence analysis was performed using MacVector version 14 (MacVector, Inc.; Apex, North Carolina) using the MHV-A59 infectious clone reference genome (GenBank accession no. AY910861). The nucleotide sequences of the amplicon and sequencing primers are available upon request. Sequencing of nsp12 and nsp14 from intermediate passages was performed using TRIzol-purified RNA from infected monolayers and using the primers listed below. Primers 6M1F (5’-TTTTGGCGAGATGGTAGC-3’) and 7M2R (5’-GGTAAGACAGTTTTAGGTGAG-3’) were used to generate a 3,425 nucleotide amplicon containing all of nsp12. Primers 7M3F (5’-ATGCTTACCAACTATGAGC-3’) and 8M3R (5’-CCGATTTGAATGGCGTAG-3’) were used to generate a 2,713 nucleotide amplicon containing all of nsp14. The PCR conditions for these reactions are the same as those used to generate the amplicons used for full-genome sequencing (16).

### Replication and RNA synthesis kinetics

Viral replication kinetics in DBT-9 cells were determined at an MOI of 1 PFU/cell or MOI of 0.01 PFU/cell as described previously (15). Supernatant (300 μL) was harvested at the indicated time points, and the virus titer was determined by plaque assay. The accumulation of genomic RNA at an MOI of 1 PFU/cell was measured by two-step real-time quantitative RT-PCR (RT-qPCR) using intracellular RNA that was TRIzol-extracted at the time points indicated. RNA was reverse-transcribed using SuperScript III (Invitrogen), and cDNA derived from intracellular positive-sense viral RNA was measured using primers directed to nsp10. Values were normalized to levels of the endogenous control glyceraldehyde-3-phosphate dehydrogenase (GAPDH). No mutations within the primer binding sites emerged in either P250 population. The primers and amplification conditions are the same as reported previously (15), except that the RT product was diluted 1:10 prior to use. Samples were plated in technical duplicate to minimize well-to-well variation. Data are presented as 2-ΔCT, where ΔCT denotes CT (target, nsp10) minus CT (reference, GAPDH).

### Determination of specific infectivity

Subconfluent monolayers of DBT-9 cells in 24-well plates were infected with the indicated virus at an MOI of 1 PFU/cell, and supernatant was harvested at 12 h.p.i. The levels of genomic RNA in supernatant were measured using one-step real-time quantitative RT-PCR (RT-qPCR) on TRIzol-extracted RNA as described previously (17). Briefly, genomic RNA was detected with a 5’ 6-carboxyfluorescein (FAM) and 3’ black hole quencher 1 (BHQ-1) labeled probe targeting nsp2 (Biosearch Technologies, Petaluma, CA), and RNA copy number was calculated by reference to an RNA standard derived from the MHV A fragment. Samples were plated in technical duplicate to minimize well-to-well variation. Titers were determined by plaque assay in DBT-9 cells, and specific infectivity was calculated as PFU per supernatant genomic RNA copy.

### Nucleoside and base analog sensitivity assays

5-azacytidine (AZC), 5-fluorouracil (5-FU), and ribavirin (RBV) were purchased from Sigma (product numbers A2385, F6627, and R9644, respectively). Stock solutions of 5-FU and RBV were prepared in dimethyl sulfoxide (DMSO). 2’-C-methyladenosine (CMeA) was received from Gilead Sciences, Inc (Foster City, CA). Sensitivity assays were performed as described previously (16), except in 24-well plates at an MOI of 0.01 PFU/cell. Supernatants were harvested at 24 hours post-infection, and titers were determined by plaque assay.

### Phyre^2^-modeling of MHV-nsp14

The MHV nsp14 structure was modeled with the Phyre^2^ online program (63) using nsp14 residues 3-519, corresponding to residues 6056-6573 of the ORF1ab polyprotein. The model was analyzed using the Pymol Molecular Graphics System, Version (Schrödinger, LLC).

### Generation of nsp12 and nsp14 swaps

Viruses containing nsp12, nsp14, or nsp12 and nsp14 from MHV-ExoN(-) P250 were generated using the MHV-A59 reverse genetics system (91). RNA from the MHV-ExoN(-) P250 virus was reversed transcribed with SuperScript III (Invitrogen) and used to generate amplicons containing either nsp12 or nsp14. Each amplicon was flanked by 15 bp that overlapped with an amplicon generated from the backbone plasmid. Amplicons were inserted into MHV-A59 fragments using the InFusion HD Cloning kit (Takara Bio USA, Inc, Mountain View, CA). Nsp12 is split across MHV E and F, while nsp14 is contained within MHV F. Reaction mixtures contained 50ng of vector, 200ng of insert, and 2μL of enzyme and were incubated for 15 minutes at 50°C. Errors were corrected by site-directed mutagenesis using Pfu Turbo polymerase (Agilent, Santa Clara, CA). The nsp12/14-P250 swap was generated through restriction digestion of the individual swaps using BsmBI and StuI followed by gel purification and assembly using T4 DNA ligase (NEB, Ipswich, MA). Viable viruses were constructed and rescued as described previously (91).

### Competitive fitness assays

Competitor viruses were competed with an MHV-ExoN(-) virus harboring 10 silent mutations in the probe-binding region within nsp2. Subconfluent DBT-9 monolayers in 24-well plates were coinfected at a total MOI of 0.01 PFU/cell with competitor and reference viruses at a 1:1 ratio and passaged 4 times. For each passage, supernatants were harvested at 24 hours. RNA was extracted from 100μL of supernatant using 900μL of TRIzol reagent and PureLink RNA Mini Kit columns (Thermo Scientific, Waltham, MA), and 150μL was used to infect fresh cells in a 24-well plate (total MOI estimated at 1 PFU/cell). The proportion of each virus was determined by real-time RT-qPCR from the infection supernatant using two Taqman probes with different fluorescent dyes in separate reactions. Competitor viruses were detected with the same probe used in specific infectivity analyses (14). Reference viruses were detected by a probe targeting the same region but with 10 silent mutations (5’-TCCGAACTACTGCAACCCCAAGTG-3’) and labeled with 5’ Quasar 670 and 3’ black hole quencher 2 (BHQ-2) (Biosearch Technologies, Petaluma, CA). RNA copy number was calculated by reference to an RNA standard generated by *in vitro* transcription of the corresponding MHV A fragment, and relative RNA abundance was calculated as the ratio of competitor genomes to reference genomes.

### Statistical analysis

GraphPad Prism 6 (La Jolla, CA) was used to perform statistical tests. Only the comparisons shown [e.g. ns or asterisk(s)] within the figure or legend were performed. In many cases the data were normalized to untreated controls. This was performed using GraphPad Prism 6. The number of replicate samples is denoted within each figure legend.

### Accession numbers

Full-length genome sequences for WT-MHV P250 and MHV-ExoN(-) P250 have been deposited in GenBank (accession numbers MF618252 and MF618253, respectively).

## ACKNOWLEDGEMENTS

We thank members of the Denison laboratory for valuable discussions. This work was supported by United States Public Health Service awards R01-AI108197 (M.R.D), T32-GM007347 (K.W.G), F30-AI129229 (K.W.G), T32-HL07751 (J.B.C), T32-AI089554 (N.R.S.), T32-AI095202 (E.C.S.), and F32-AI108102 (E.C.S.), all from the National Institutes of Health. The content is solely the responsibility of the authors and does not necessarily represent the official views of the National Institutes of Health. The authors declare no conflicts of interest.

---

**Supplemental Figure 1. Deleted regions within WT-MHV P250 and MHV-ExoN(-) P250.**

